# Gut microbial conversion of dietary elderberry extract to hydrocinnamic acid improves obesity-associated metabolic disorders

**DOI:** 10.1101/2025.05.18.654739

**Authors:** Sara Alqudah, Beckey DeLucia, Lucas J. Osborn, Rachel L. Markley, Viharika Bobba, Sarah M. Preston, Tharika Thambidurai, Layan Hamidi Nia, Clifton G. Fulmer, Naseer Sangwan, Ina Nemet, Jan Claesen

## Abstract

Obesity-associated metabolic disorders such as type 2 diabetes mellitus and metabolic dysfunction associated fatty liver disease are major global health concerns, yet current pharmacological treatments often present with major side-effects. Dietary interventions including polyphenol-rich foods offer a promising complementary option for obesity amelioration, but their efficacy is dependent on specific gut microbial metabolism and the underlying molecular mechanisms mostly remain elusive. Here, we demonstrated that dietary elderberry (Eld) extract abrogates the effects of an obesogenic diet in a gut microbiota-dependent manner, by preventing insulin resistance and reducing hepatic steatosis in mice. We developed a targeted, quantitative liquid chromatography-tandem mass spectrometry method for detection of gut bacterial polyphenol catabolites and identified hydrocinnamic acid as a key microbial metabolite, enriched in the portal vein plasma of Eld supplemented animals. Next, we showed that hydrocinnamic acid potently activates hepatic AMP-activated protein kinase α, explaining its role in improved liver lipid homeostasis. Furthermore, we uncovered the metabolic pathway cumulating in hydrocinnamic acid production in the common gut commensal *Clostridium sporogenes*. Our characterization of hydrocinnamic acid as a diet-derived, and microbiota-dependent metabolite with insulin-sensitizing and anti-steatotic activities will contribute to microbiome-targeted dietary interventions to prevent and treat obesity-associated metabolic diseases.

## Introduction

Obesity-associated metabolic disorders continue to increase in both incidence and severity, posing a challenge for our healthcare system^1^. Obesity initiates a sequence of metabolic disturbances, beginning with impaired glucose metabolism, which leads to hyperglycemia and subsequent hyperinsulinemia, the hallmark indicators of prediabetes^2^. If left untreated, prediabetes progresses with the development of insulin resistance due to the loss of a glucose-stimulated acute insulin response. The resulting increased demand on insulin-producing pancreatic β-cells results in their dysfunction, eventually developing into type 2 diabetes mellitus (T2DM)^3^. Furthermore, obesity is closely associated with an elevated risk of metabolic dysfunction-associated fatty liver disease (MAFLD), primarily due to an imbalance between hepatic fatty acid uptake from plasma and de novo fatty acid synthesis, as well as fatty acid export as triglycerides within very-low-density lipoproteins (VLDL). This imbalance leads to lipid accumulation in hepatocytes, contributing to hepatic steatosis^4^.

Sex significantly influences several aspects of susceptibility to, and progression of obesity-related metabolic diseases. Specifically, men exhibit a higher propensity for obesity compared to women, which in part can be explained by enhanced adiposity when subjected to obesogenic diets^5,6^. Moreover, men naturally exhibit higher insulin resistance, due to lower Hexokinase II expression, which decreases glucose uptake into skeletal muscle cells ^7,8^. Finally, premenopausal women have a lower prevalence of MAFLD, attributed to the protective effects of estrogen^9,10^.

The AMP-activated protein kinase (AMPK) signaling pathway provides one of the key molecular targets for obesity treatment. AMPK functions as a cellular energy sensor, activated in response to reduced intracellular ATP levels, and plays a central role in maintaining energy homeostasis across all eukaryotic cells. Activation of AMPK through the phosphorylation of the α subunit, has been shown to enhance glucose uptake, promote fatty acid oxidation, and stimulate mitochondrial biogenesis^11–13^. In addition, AMPK activation suppresses lipogenesis and cholesterol biosynthesis^14,15^. In the context of metabolic disorders, dysregulation of AMPK signaling has been consistently observed^16–18^. Metformin and lifestyle changes are the first-line treatment of obesity-induced T2DM^19^. Although metformin decreases blood glucose levels by a hepatic AMPK-dependent mechanism, it has undesirable side effects, including upper respiratory infection, abdominal distension, dizziness, and vomiting^20^. Sulfonylureas, GLP-1 receptor agonists, and thiazolidinediones are also used to treat T2DM, but can present with even more severe side effects such as low glucose levels, pancreatitis, bone loss, hepatotoxicity, and heart failure^21^.

Lifestyle changes including improved exercise regimes and dietary interventions provide alternative options for the treatment and prevention of metabolic diseases. Colorful fruits and vegetables are an important component of a healthy diet and they are rich in polyphenols, including flavonoids and other phenolic acids^22–24^. Despite various health benefits being attributed to polyphenol consumption, key mechanistic insights remain to be discovered as they do not reach substantial systemic concentration after ingestion in the diet or as a supplement. The enzymatic repertoire expressed by the human host is not capable of extensive polyphenol metabolism, leading to the relatively unscathed passage of these compounds throughout the gastrointestinal tract. However, there is a growing interest in the metabolic transformations catalyzed by select members of the gut microbiota, resulting in smaller bioactive metabolites that can be absorbed in the colonic epithelium and enter the bloodstream^24–27^.

We previously reported that the anti-steatotic effects of dietary interventions with various berry extracts as the primary source of polyphenols hinge on gut microbial metabolism^28^. In particular, elderberry (Eld; *Sambucus nigra*) is a rich source of anthocyanins, flavanols, as well as other phenolic acids^29^. We showed that the gut microbial end-product 4-hydroxyphenylacetic acid (4-HPAA), a monophenolic acid (MPA), was sufficient to reverse obesity-associated fatty liver disease in male mice^28^. In the current work, we hypothesized that polyphenol-supplemented diets require gut microbial MPA production for improving host metabolic parameters in prediabetes and MAFLD development. We used Eld as a source of polyphenols, and protocatechuic acid (PCA), a bioactive MPA, as a positive control. Additionally to addressing the role of the gut microbiota in metabolic diseases, we assessed sex-specific differences in both male and female mice. This revealed an improvement in insulin homeostasis in males and females, and a prevention of hepatic steatosis in males only. Using LC-MS/MS analysis, we identified hydrocinnamic acid (or 3-phenylpropionic acid) as a potent AMPKα agonist and demonstrated that this MPA can be formed by a gut commensal *C. sporogenes* from Eld extract. Our work highlights a gut microbial pathway that can be stimulated with a prebiotic intervention, contributing to the management of prediabetic and MAFLD associated pathology.

## Results

### Dietary supplementation with elderberry extract reduces high fat diet-induced weight gain

To assess the impact of dietary polyphenols in the context of improving obesity-associated metabolic parameters, we used a high fat diet (HFD)-induced obesity model, in which 4-week-old mice were randomly assigned to dietary groups with ad libitum access to a HFD (negative control group) or a HFD supplemented with 1% elderberry (Eld) extract as a polyphenol source. An additional positive control group received a HFD supplemented with 1% protocatechuic acid (PCA), a well-known monophenolic acid (MPA) formed from gut bacterial polyphenol catabolism involved in modulation of metabolic disease parameters^24^. We analyzed both male and female mice to account for metabolic sex-differences. To address the contribution of the microbiome, we compared the aforementioned groups on regular drinking water to animals that received an antibiotic cocktail (ABX; 1 g/L neomycin, 1 g/L ampicillin and 0.5 g/L vancomycin) during the course of the 12-week experiment.

We measured body weights at weekly intervals and as expected found that male HFD control mice had the highest weight gain (Fig. 1A and B). Male weight gain was attenuated in both the Eld and PCA supplemented groups, tracking with our observations in female mice (Fig. 1C and D). These observed differences were not due to animal aversion to the flavor of the dietary additives, as both male and female mice had higher food intake on the Eld or PCA supplemented HFD diets (Fig. 1E-H). Gut microbiome depletion in HFD mice led to a decrease in weight gain, but did not significantly impact body weight in Eld or PCA supplemented animals. No significant differences were observed in overall body composition (proportions of fat mass and lean mass to total body mass) among the experimental groups in either males or females (SI Fig. S1A-S1D).

**Fig. 1.**
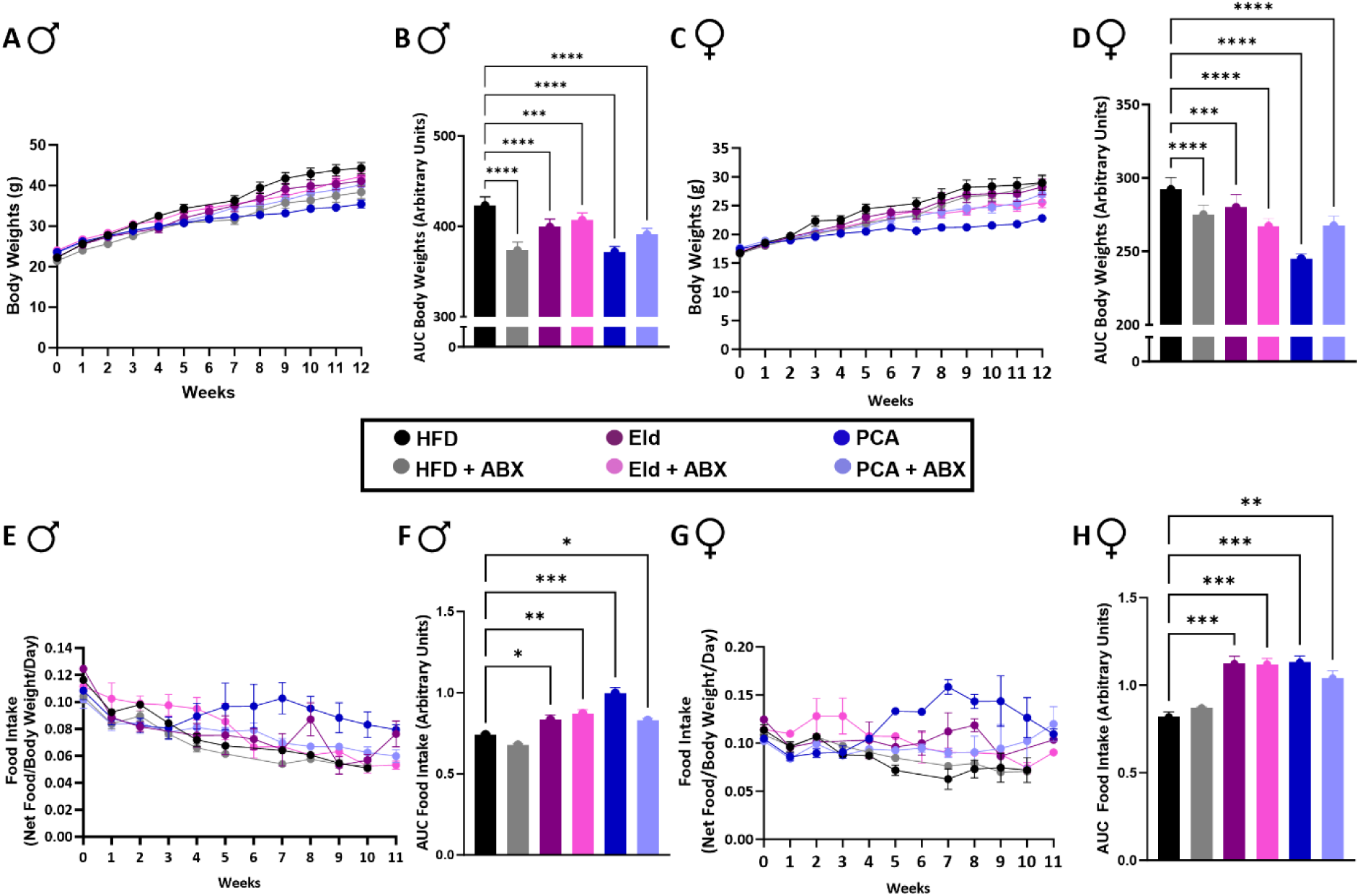
Elderberry extract reduces high fat diet-induced weight gain despite an increase in food intake. **(A, C)** Body weight (g) of 4-week (wk)-old male (A) and female (C) C57BL/6 mice fed a control high fat diet (HFD), or the same diet supplemented with 1% w/w elderberry extract (Eld), or 1% w/w protocatechuic acid (PCA), with regular or antibiotic (ABX) drinking water for 12 wks; n = 9–10 per group. **(B, D)** Mean cumulative area under the curve (AUC) for body weights after 12 wks. **(E)** Food intake (net food/weight/day) for all male groups after 12 wks of diet, **(F)** Mean cumulative area under the curve (AUC) for food intake after 12 wks. **(G)** Food intake (net food/weight/day) for all female mice groups after 12 wks of diet. **(H)** Mean cumulative area under the curve (AUC) for food intake after 12 wks. Error bars represent SEM. Statistical analysis was performed via ANOVA.

### Eld supplementation improves insulin sensitivity in a microbiome-dependent manner

To evaluate glucose metabolism and homeostasis, we performed a glucose tolerance test (GTT) following a four hour fast on week 10 of the study. In male animals with an intact microbiome, blood glucose clearance was significantly improved in both the Eld and PCA groups compared to the HFD control (Fig. 2A and SI Fig. S2A), though this trend was not observed in female mice (Fig. 2C and SI Fig. S2B). This indicates that dietary supplementation with either parent polyphenols or their microbial metabolites can improve glucose metabolism in a sex-dependent manner. These sex differences in glucose tolerance and fasting blood glucose levels could be attributed to female mice generally exhibiting better glucose tolerance and being more refractory to obesity development^5,30^. The groups of mice that had their gut microbiota depleted by ABX overall showed a similar improvement in glucose clearance and fasting blood glucose, irrespective of their diets (Fig. 2E and 2F).

**Fig. 2.**
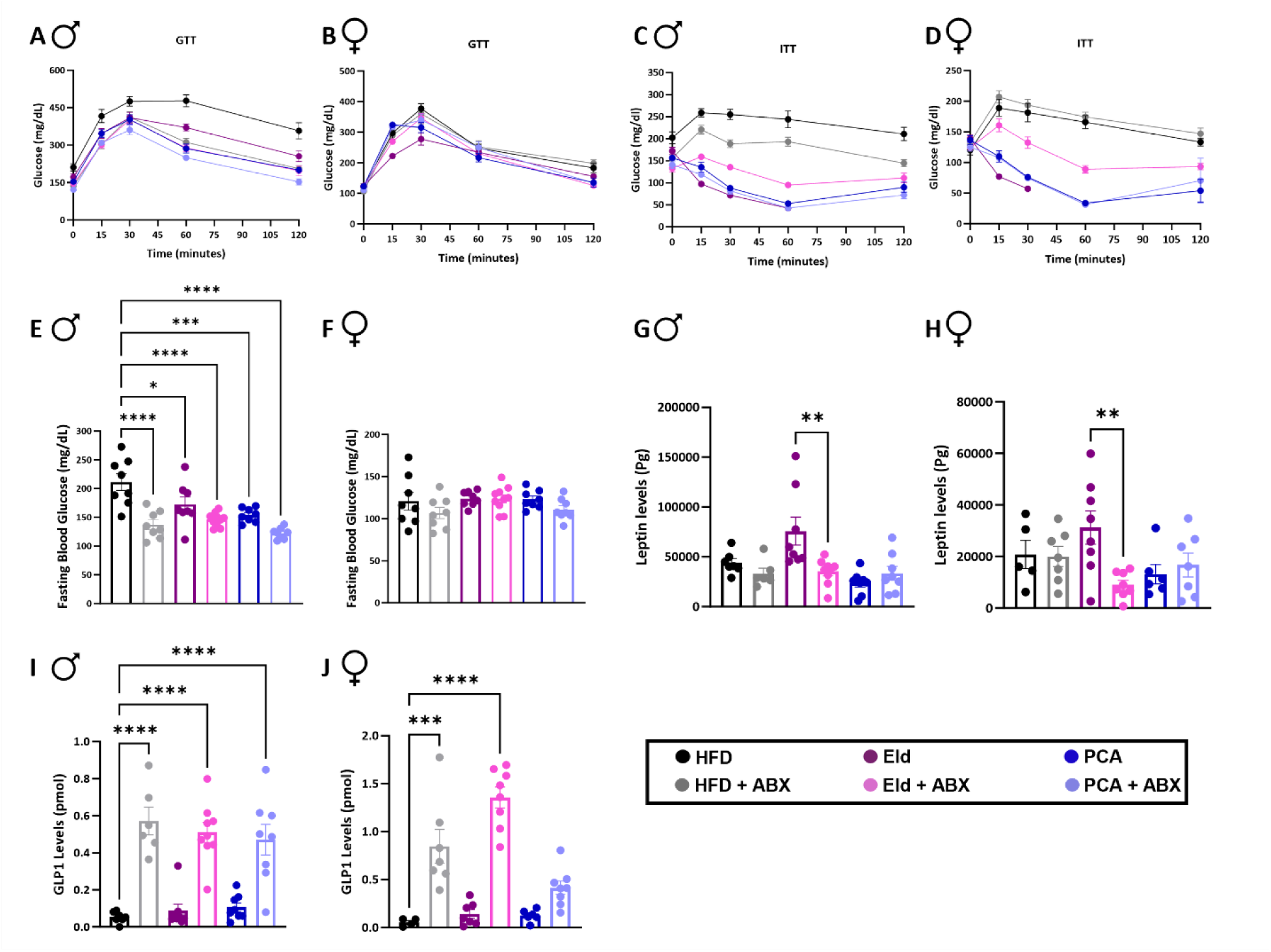
Elderberry extract substantially increased insulin sensitivity, glucose homeostasis, and leptin levels. **(A-B)** Glucose tolerance test (GTT) for all male and female mice groups on wk 10 following a 4 hours fast, n = 9–10 per group. **(C-D)** Insulin tolerance test (ITT) (mg/dL) for male and female mice groups on wk 11 following a 4 hours fast, n = 9–10 per group. **(E-F)** Fasting blood glucose for male and female mice following 4 hours fasting. End point male and female plasma **(G-H)** leptin, **(I-J)** GLP1. Error bars represent SEM. Statistical analysis was performed via one-way ANOVA.

One week after the GTT (11 weeks on diet), we performed insulin tolerance tests (ITT) to assess insulin sensitivity in our animals. Male mice supplemented with either Eld or PCA had significantly lower insulin resistance compared to the control animals (Fig. 2C and SI Fig. S2C). Interestingly, the effect of Eld was in part microbiome-dependent, as the Eld-ABX animals had a blood glucose response intermediate between the Eld and HFD groups. This emphasizes the importance of gut microbial metabolism in obtaining health benefits from Eld polyphenol supplementation. As anticipated, the microbiota did not have a noticeable effect on PCA supplementation. The improved insulin tolerance in Eld and PCA groups and importance of the gut microbiome for Eld metabolism replicated in female mice (Fig. 2D and SI Fig. S2D). It is of note that because of the profound insulin sensitivity effect, animals in the Eld group received glucose injections midway through the experiment to alleviate the discomfort. Overall, our findings suggest that the gut microbiota are required for metabolizing elderberry extract into bioactive metabolites that improve insulin tolerance in both male and female mice.

To further profile our animals’ metabolic state, we used a multiplex approach to measure the endpoint plasma concentration of gut active hormones. Interestingly, Male and female leptin levels in Eld treated animals were significantly higher compared to their Eld-ABX counterparts, indicating that a microbial byproduct of polyphenol metabolism may boost leptin production (Fig. 2G,H). Leptin is produced by adipose tissue and is involved in regulating energy balance by limiting appetite^31^. This finding is particularly interesting since we did not observe differences in fat mass (SI Fig. S1A-S1D). Glucagon-like peptide 1 (GLP-1) levels were overall higher across ABX treated groups, irrespective of their diets (Fig. 2I,J). We did not observe differences in the levels of insulin, glucagon, and peptide YY among the treatment groups (SI Fig. S2E-2J).

### Eld supplementation requires an intact gut microbiome for hepatic steatosis prevention in male mice

Here, we assess the contribution of the microbiota to Eld metabolism in male and female animals. Hematoxylin and eosin (H&E)-stained liver histology slides were prepared and evaluated in a blinded manner by a board-certified pathologist. Liver sections from male Eld or PCA groups had reduced lipid accumulation compared to the HFD controls (Fig. 3A). The animals on Eld-ABX did not have a reduction in hepatic lipid droplet accumulation, emphasizing the importance of gut microbial metabolism in mediating the health benefits of dietary polyphenols. As expected, the steatosis protective effect of the monophenolic acid PCA is not significantly affected by the absence of the gut microbiota. Most notably, mice supplemented with Eld showed significant improvements in steatosis percentage (Fig. 3B), steatosis score (Fig. 3C), NAFLD activity score (Fig. 3D), and inflammation score (Fig. 3E) compared to their counterparts on Eld-ABX with an ablated gut microbiome. The female control mice only had minimal hepatic lipid accumulation after 12 weeks on HFD, which precluded us from further analyzing the histological effects of Eld, PCA or the microbiota on their livers (SI Fig. S3). This is consistent with previous studies showing that female mice are naturally more resistant to HFD-induced metabolic changes^7,30^. Our finding that ABX ablation impedes the effects of an Eld-supplemented diet further supports the gut microbiome’s role in protecting against steatosis in obese mice.

**Fig. 3.**
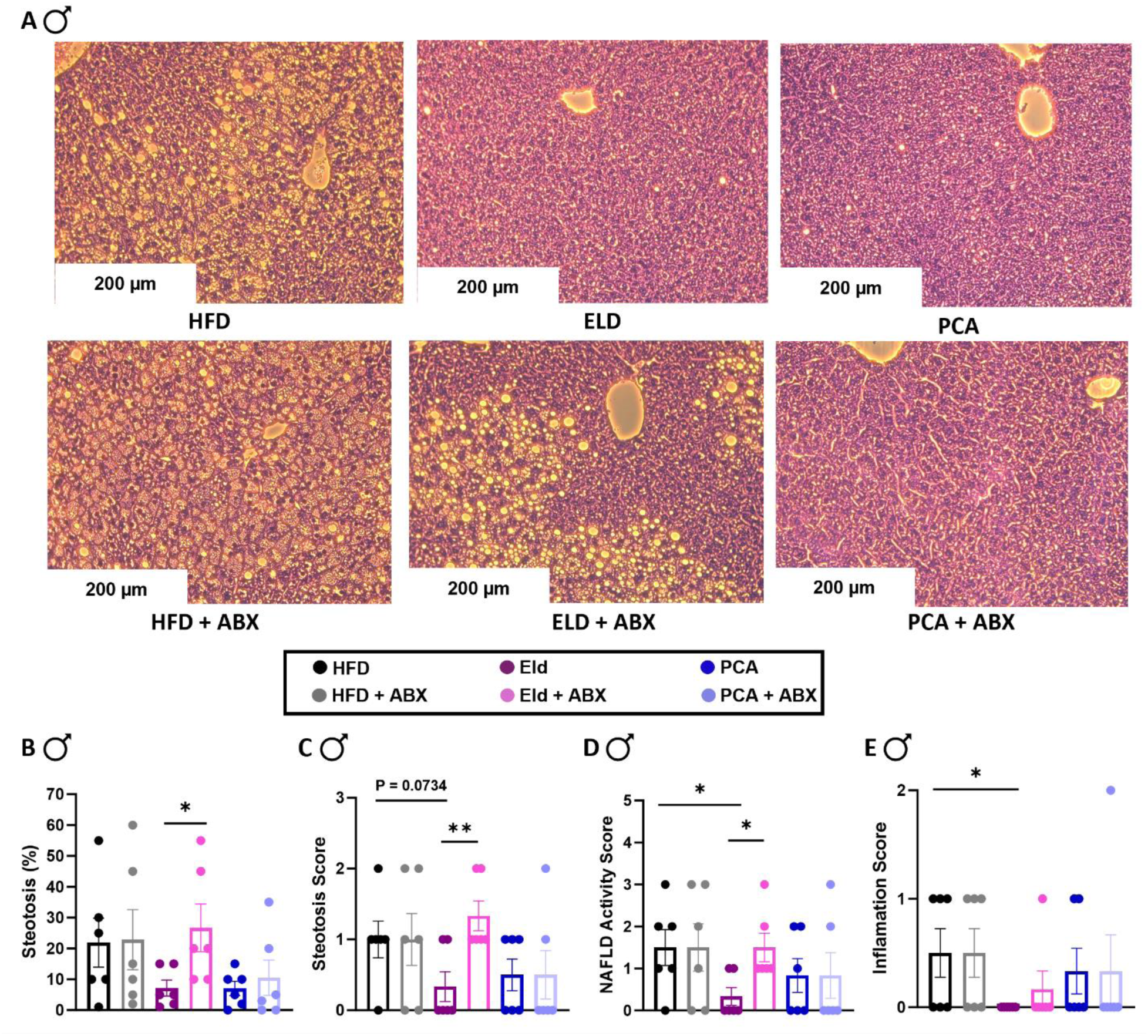
Elderberry extract prevents hepatic steatosis in male mice in a microbiome-dependent manner. **(A)** After 12 wks of the different diets, H&E-stained sections of the male mouse livers. Representative images shown from n = 6 per group. Pathologic assessment for the H&E-stained sections **(B)** Steatosis percentage **(C)** Steatosis score **(D)** NAFLD activity score **(E)** Inflammation. Error bars represent SEM. Statistical analysis was performed using unpaired two-tailed Student’s t test.

### Eld mice have increased hepatic AMPKα activation and improved lipid metabolism

We tested for hepatic AMPKα activation via increased T172 phosphorylation by immunoblot analysis and found that both male and female mice showed increased AMPKα phosphorylation on Eld, but only when the gut microbiome remained intact (Fig. 4A and 4B, SI Fig. S4C and S4D). No differential activation was observed between the HFD and HFD-ABX control groups (Fig. 4A and 4B, SI Fig. S4E and S4F). These findings suggest that hepatic AMPKα activation may be in part responsible for the steatosis prevention observed in the livers of males on Eld (Fig. 3A) and further support the importance of the gut microbiome in this process. As expected, PCA-induced AMPKα activation (Fig. 4A and 4B, SI Fig. S4G-S4J) is not dependent on the microbiota, which is consistent with previous studies on PCA in mice^32–34^. We next analyzed phosphorylation of acetyl-CoA carboxylase (ACC), the enzyme catalyzing the committing step to fatty acid biosynthesis, and a downstream target of AMPKα regulation. However, no significant Eld-dependent differences were observed (SI Fig. S5A and S5B).

**Fig. 4.**
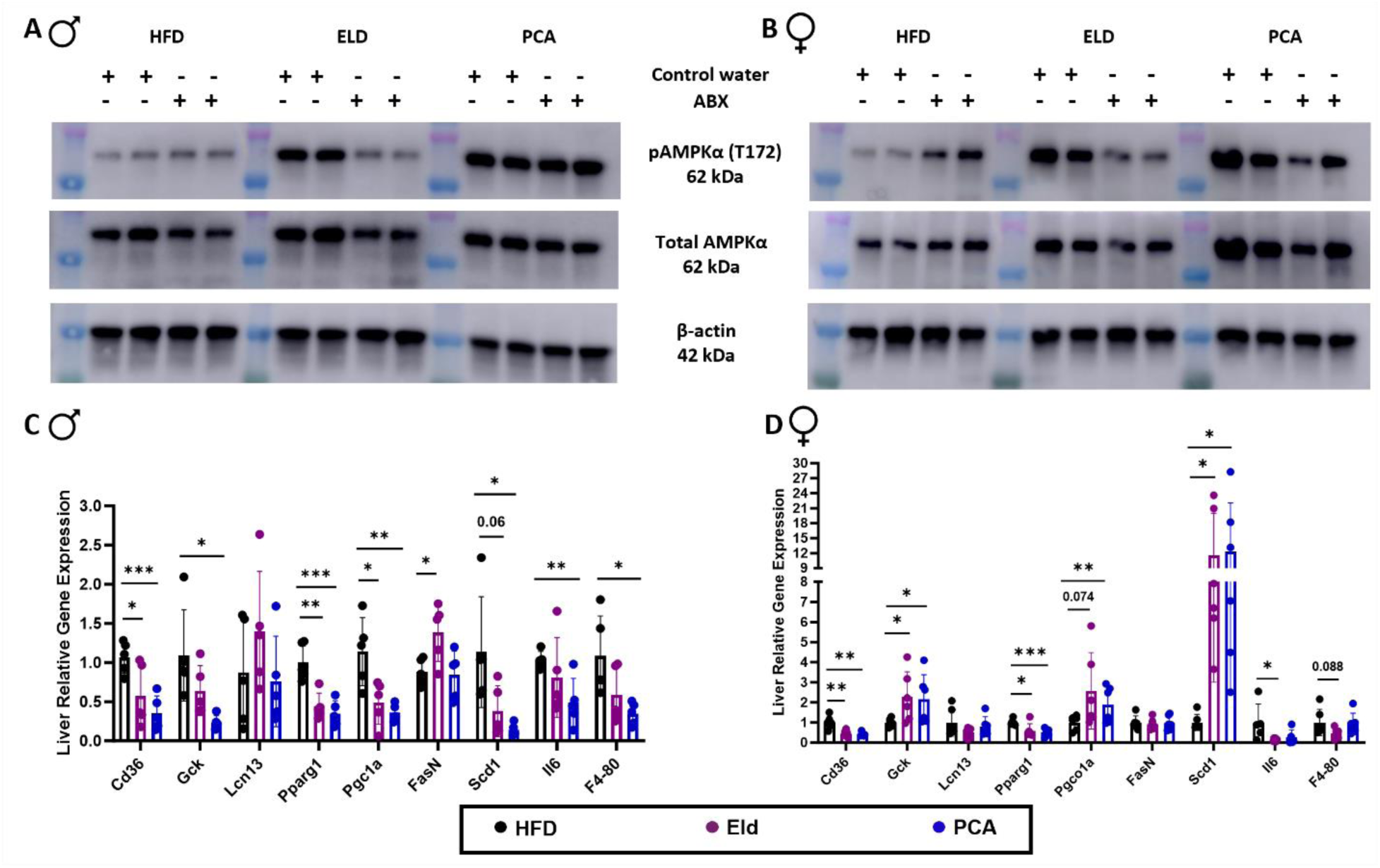
Elderberry extract increases the activation of AMPKα and impacts hepatic gene expression in male and female mice in a microbiome-dependent manner. **(A-B)** Western blot analysis of pAMPKα (T172), total AMPKα, and β-actin in male and female mice after 12 wks of the different diets and water**. (C-D)** RT-qPCR quantification of mRNA transcripts involved in metabolism and inflammation of the male and female mice liver normalized to CycloA n=6 per group. Error bars represent SEM. Statistical analysis was performed using unpaired two-tailed Student’s t test.

To further investigate metabolic changes, mRNA expression of several liver genes linked to lipid metabolism and inflammation were measured (Fig. 4C and 4D). In both male and female mice supplemented with Eld or PCA, transcripts of the fatty acid transporter *Cd36*, and peroxisome proliferator-activated receptor gamma (*Pparg1*), a key regulator of adipogenesis, lipid metabolism, and insulin sensitivity, were markedly reduced. Some genes showed sex-specific differences, including glucokinase (*Gck*) and peroxisome proliferator-activated receptor gamma coactivator-1 alpha (*Pgc1a*), a master regulator of mitochondrial biogenesis. Notably, stearoyl-CoA desaturase 1 (*Scd1*), which converts saturated fatty acids into monounsaturated fatty acids, showed opposing patterns between sexes. In male mice, *Scd1* expression was significantly reduced in both the Eld and PCA groups, while in female mice, expression levels were higher compared to controls. This finding agrees with previous reports showing that *Scd1* is more highly expressed in female skeletal muscle and adipose tissue due to sex differences in fatty acid metabolism^8^. Our study now also reports this difference in liver tissue (Fig. 4D). Taken together, these results suggest that both Eld metabolites and PCA promote monounsaturated fatty acid synthesis in the liver of female mice while reducing it in males. Additionally, an Eld diet lowered gene expression of the liver inflammation markers IL-6 and F4-80 in female mice only (Fig. 4D).

### Eld fed mice cecal microbiomes are enriched in polyphenol metabolizing commensals

To investigate a potential link between the observed sex differences and the gut microbiota, and to identify potential polyphenol catabolizing microbes, we determined the cecal microbial composition using 16S rRNA sequencing. Overall, control and PCA female animals had a higher alpha-diversity compared to males on the same diet as measured by Simpson index (Fig. 5A). While the alpha-diversity of the Eld male and female mice did not differ significantly, their community compositions distinctly clustered separate from each other, as well as from the other groups (Fig. 5B). These differences are also reflected in the total diversity plots (Fig. 5C) and heat map (Fig. 5D), which highlight an overall increased abundance of *Faecalibaculum* in males, as well as treatment-specific sex-differences, most notably *Lactobacillus*, *Lachnospiraceae* from the NK4A136 group, and *Roseburia*. Members of these bacterial genera are linked to improvements in hepatic steatosis and inflammation through the production of short-chain fatty acids^35–38^. Additionally, female mice on Eld displayed an increased abundance of *Akkermansia* and *Bacteroides*, bacterial taxa previously implicated in beneficial metabolic effects^39,40^. Bacterial genera that are typically associated with polyphenol metabolism, including *Lachnospiraceae* and *Lactobacillus*, were among the top contributors to the observed treatment-dependent beta-diversity in both male and female animals (Fig. 5F).

**Fig. 5.**
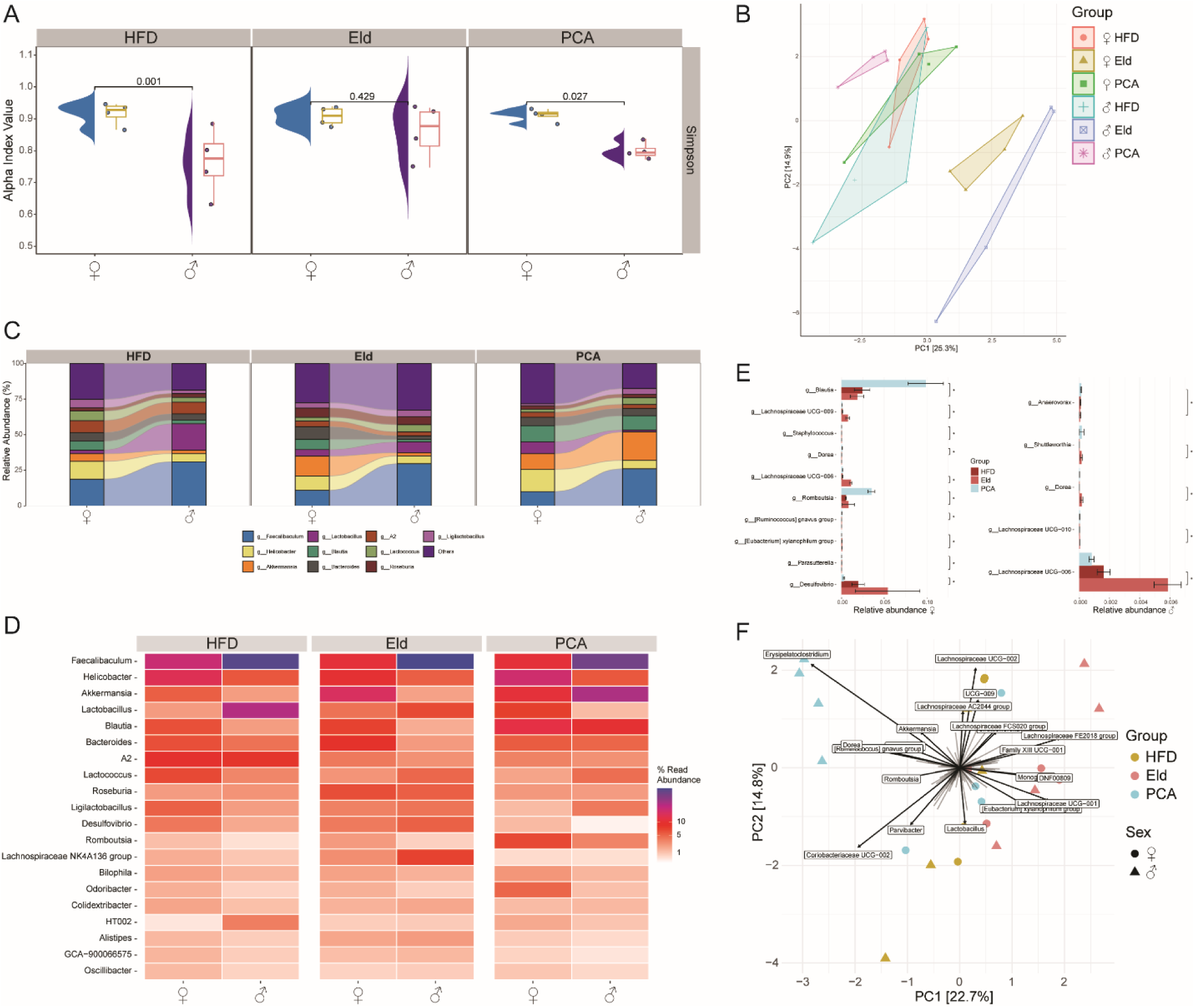
Eld mice have an increased relative abundance of health-associated gut bacteria. **(A)** Simpson alpha diversity estimates compared between male and female cecal communities on the different diets. Pairwise analysis was done using Wilcoxon rank sum t-test. **(B)** Principal component analysis (PCA) plot based on the Bray–Curtis dissimilarity index between the cecal 16S rRNA profile of all the six groups. **(C)** Stacked bar plot comparison of male and female total diversity by diet, representing the most abundant taxa in each group. **(D)** Heatmap of the differentially abundant bacterial taxa across comparing male and female microbiota across the different diet groups. **(E)** Differential abundance analysis plots for statistically significant taxa in the male (left) and female (right) cecal communities across the different diet groups. Pairwise test used was “metagenomeSeq” with Benjamini Hochberg correction. **(F)** Principal component analysis plot showing the Bray-Curtis dissimilarity between groups and highlight bacterial taxa that significantly contribute to beta diversity. (n = 4 animals for all 16S rRNA sequencing analyses).

### Polyphenol-derived microbial catabolites are increased in the portal plasma from animals on Eld supplemented diet

To gain functional insights into which bioactive microbial metabolites contribute to the beneficial effects of Eld supplementation, we developed a targeted liquid chromatography-tandem mass spectrometry (LC-MS/MS) stable isotope dilution method for measuring monophenolic acids that are potential polyphenol catabolites, chromatographically separating different isomers. We analyzed plasma from the portal vein as this contains absorbed nutrients, xenobiotics, and other compounds from the gastrointestinal tract before they are processed by the liver^41^. Among the nine microbial polyphenol catabolites monitored in our LC-MS/MS method, hydrocinnamic (or 3-phenylpropionic acid) (Fig. 6A, 6B) and 3-hydroxyphenylacetic acid (3-HPAA) (Fig. 6C, 6D) were significantly elevated in both female and male mice on Eld. The Eld+ABX group did not have a similar increased portal plasma level in these MPAs. In contrast, ferulic acid (4-hydroxy-3-methoxycinnamic acid) a compound present in the Eld extract, was significantly reduced in the Eld compared to the Eld-ABX group, suggesting consumption by the microbiota (Fig. 6E, 6F). Likewise, phloretic acid (3-(4-hydroxyphenyl)propionic acid) levels were significantly higher in Eld-ABX female mice compared to their Eld counterparts (Fig. 6H). Other monophenolic acids measured showed no significant differences across groups (SI Fig. S6A, S6D) including the portal plasma levels of PCA in PCA treated animals when compared to the HFD group, suggesting potential microbial consumption of this MPA. Increased PCA levels in PCA-ABX mice compared the PCA group further corroborates the potential role of the gut microbiota in downstream PCA catabolism (Fig. 6I and 6J). It is tempting to speculate that this dietary PCA got converted to vanillic acid (4-hydroxy-3-methoxybenzoic acid) through 3-methylation in our male animals (Fig. 6K), a process which is known to be catalyzed by gut bacteria^42,43^. Intriguingly, we observed the opposite pattern in the female animals on PCA +/- ABX (Fig. 6L).

**Fig. 6.**
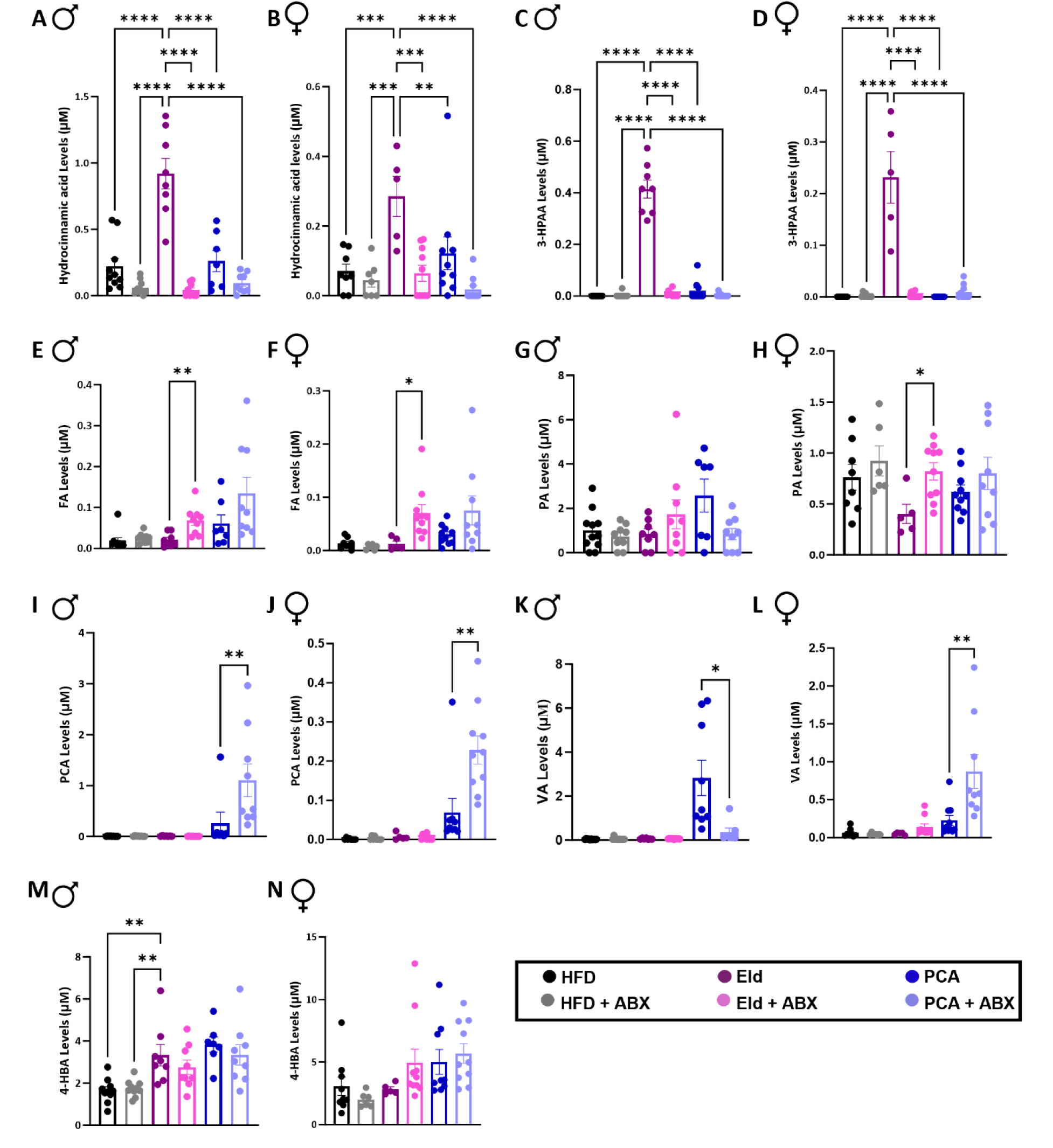
LC-MS/MS portal plasma analysis of Eld fed mice revealed an increase in 3-PPA and 3-HPAA microbial polyphenol catabolites. LC-MS/MS measured Portal plasma from male and female mice with different gut microbial metabolite n = 5–9 per group. **(A-B)** Hydrocinnamic acid. **(C-D)** 3-Hydroxyphenylacetic acid (3-HPAA). **(E-F)** Ferulic acid (FA). **(G-H)** 4-Hydroxybenzoic acid (4-HBA). **(I-J)** Protocatechuic acid (PCA). **(K-L)** Vanillic acid (VA). **(M-N)** Phloretic acid (PA). Error bars represent SEM. Statistical analysis was performed with one-way ANOVA.

### Hydrocinnamic acid activates hepatic AMPKα signaling

Since both hydrocinnamic acid and 3-HPAA were significantly enriched in the Eld male and female mice in a microbiome-dependent manner (Fig. 6A-D), we next tested their capability to activate AMPKα. *In vitro* cultivated hepatocyte-derived cellular carcinoma cells (HuH-7) were treated with either MPA. Cells treated with hydrocinnamic acid demonstrated a significant increase in AMPKα phosphorylation compared to the vehicle control (Fig. 7A). Notably, 3-HPAA treatment had no effect on AMPKα total levels nor on phosphorylation. Hence, we speculate that the AMPKα activation observed in the livers of our male and female mice (Fig. 4A and 4B) can most likely be attributed to hydrocinnamic acid. To assess the dosage of hydrocinnamic acid required to activate AMPKα, we administered the compound to HuH-7 cells in a physiologically relevant concentration range (0.25-5 μM). Even at concentrations below the portal plasma levels (Fig. 6A and 6B), hydrocinnamic acid treatment resulted in robust AMPKα phosphorylation in a dose-independent manner (Fig. 7B and 7C). Accordingly, we observed an increase in phosphorylation of the downstream enzyme ACC (SI Fig. S7A and S7B). These findings demonstrate the potent effect of hydrocinnamic acid on AMPKα activation and downstream signaling and can explain the observed *in vivo* steatosis protective effect via promotion of fatty acid oxidation while suppressing *de novo* lipogenesis in the liver.

**Fig. 7.**
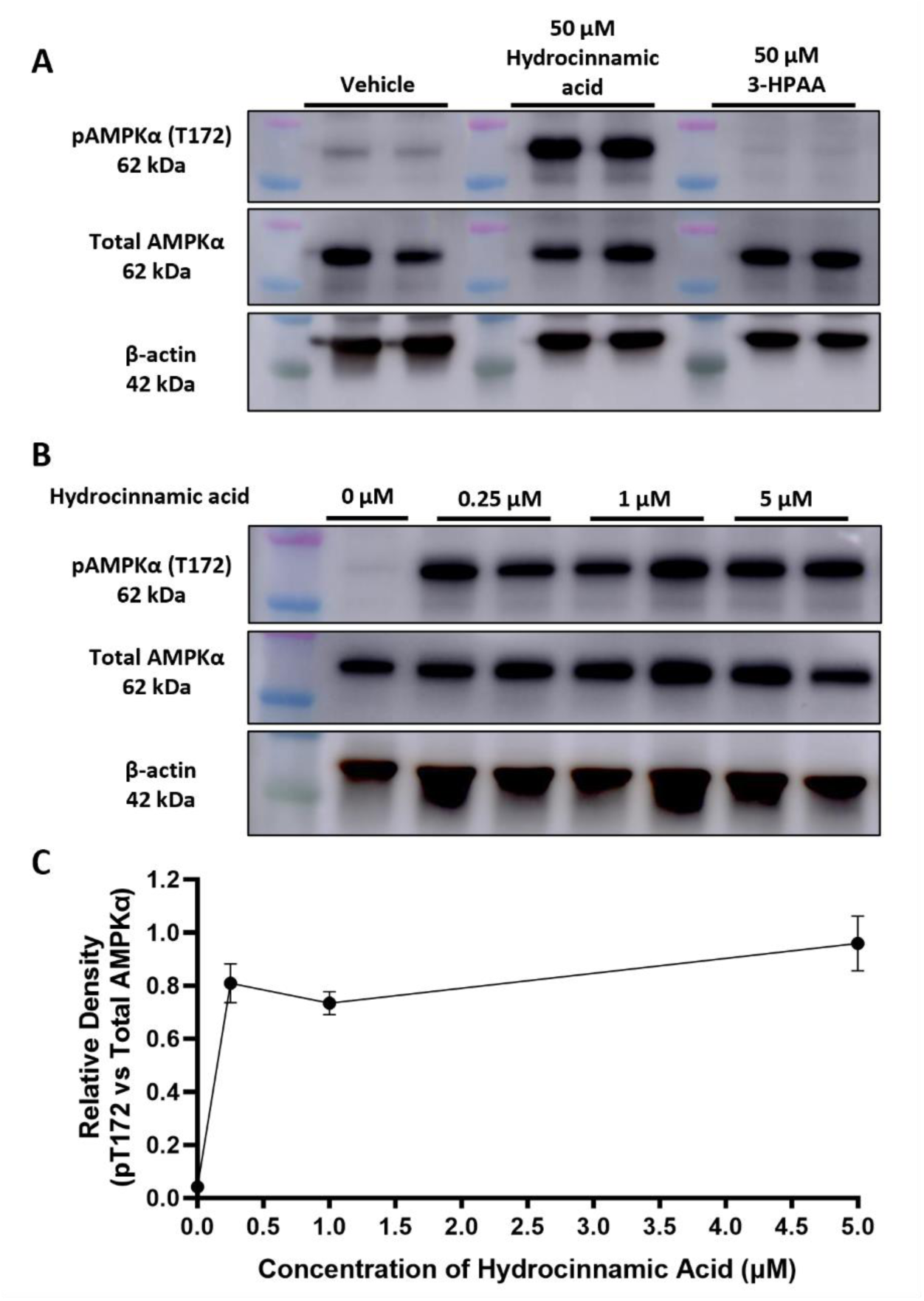
Hydrocinnamic acid activates hepatic AMPKα signaling in HuH-7 cells at physiologically relevant concentrations. **(A)** Western blot analysis of pAMPKα (T172), total pAMPKα, and β-actin for HuH-7 cells after treatment with either DMSO vehicle, 50 μM hydrocinnamic acid, or 50 μM 3-HPAA. **(B)** Western blot analysis of pAMPKα (T172), total pAMPKα, and β-actin for HuH-7 cells after treatment with DMSO vehicle control (0 μM) or 0.25, 1, and 5 μM hydrocinnamic acid. **(C)** Relative expression of the ratio of AMPKα activation after treatment with different physiological concentrations of hydrocinnamic acid. Error bars represent SEM.

### The gut commensal *Clostridium sporogenes* converts cinnamic acid from Eld into hydrocinnamic acid

Various gut bacteria are capable of hydrocinnamic acid production, including strains of *Clostridium sporogenes* and *Bacteroides fragilis*^44,45^. A main metabolic pathway for this process involves a series of enzymatic conversions starting from the aromatic amino acid phenylalanine (Phe), which has been fully characterized in *C. sporogenes* (Fig. 8A)^45^. To investigate whether these bacteria are also capable of producing hydrocinnamic acid from substrates in the Eld extract, we supplemented liquid cultures of *C. sporogenes* and *B. fragilis* with Eld extract or vehicle control under anaerobic conditions. After 4 hours of incubation, we quantified hydrocinnamic acid levels using our LC-MS/MS panel. Hydrocinnamic acid levels were higher in vehicle-treated *C. sporogenes* ATCC 15579 as compared to the no bacteria control, a baseline level which is likely due to the breakdown of Phe from the culture medium (Fig. 8B). Addition of Eld extract resulted in a significant increase in hydrocinnamic acid levels, indicating that *C. sporogenes* ATCC 15579 can additionally metabolize Eld-derived compounds into hydrocinnamic acid. In contrast, *B. fragilis* 3_1_12 HM-20 did not produce hydrocinnamic acid from either Phe present in the growth medium or Eld extract, suggesting that this strain’s metabolic capabilities differ from previously studied *B. fragilis* strains.

**Fig. 8.**
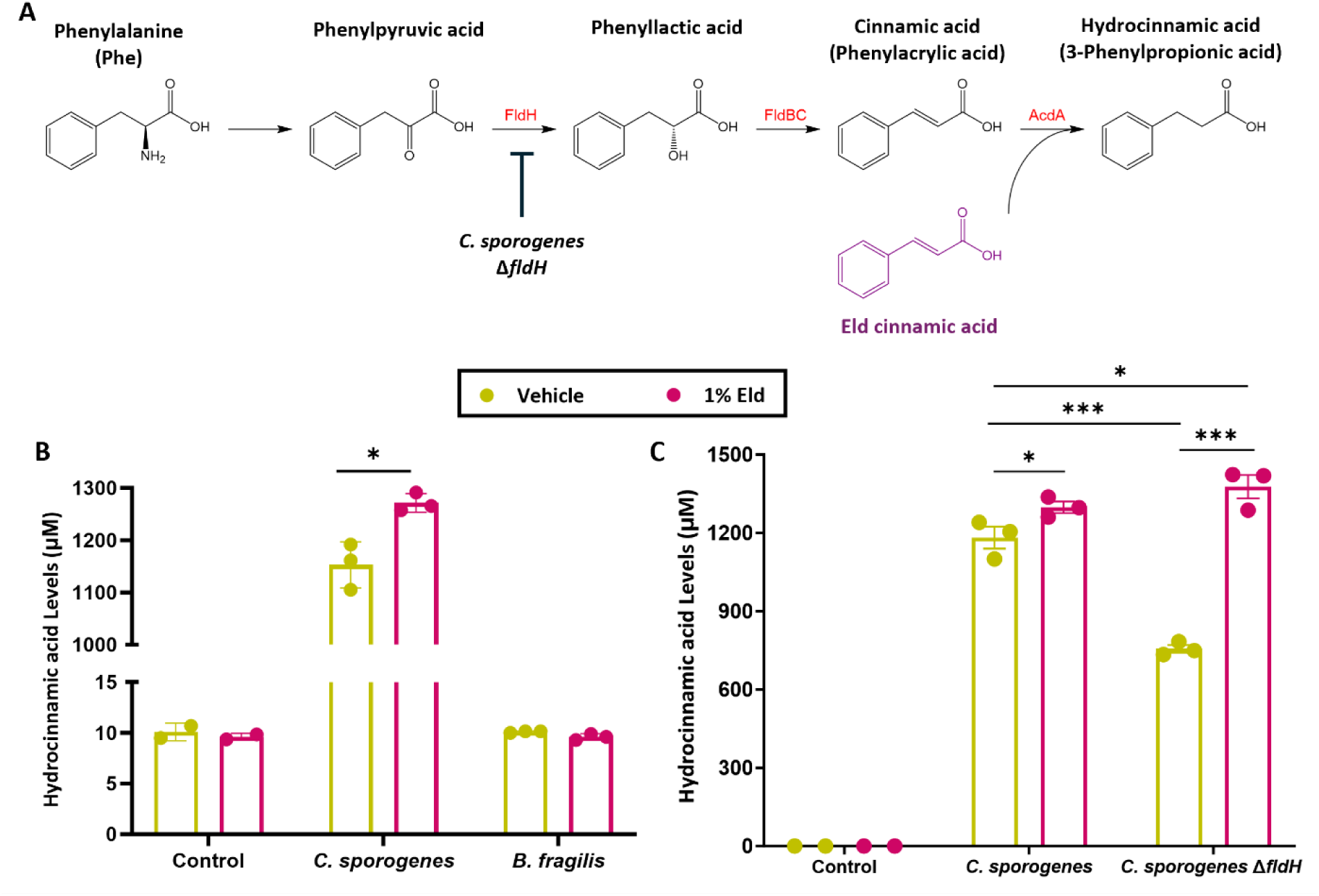
*C. sporogenes* converts cinnamic acid from elderberry extracts into hydrocinnamic acid. **(A)** The reductive pathways for phenylalanine metabolism in *C. sporogenes* to hydrocinnamic acid, with the addition of cinnamic acid from elderberry extracts. **(B)** *C. sporogenes* can convert 1% elderberry extract into hydrocinnamic acid from cinnamic acid as measured by LC-MS/MS but not *B.fragillis*, n=3 per group. **(C)** *ΔfldH C. sporogenes* can still produce the same levels of hydrocinnamic acid from elderberry extract compared to the wild-type *C. sporogenes*, as measured by LC-MS/MS, n=3 per group. Error bars represent SEM. Statistical analysis was performed using unpaired two-tailed Student’s t test.

We noticed that the Eld extract contains a substantial amount of cinnamic acid (phenylacrylic acid). Cinnamic acid can get converted to hydrocinnamic acid by 3- (aryl)acrylate reductase, the final enzymatic step in the Phe metabolic pathway (Fig. 8A), and we postulated that the Eld cinnamic acid is responsible for the increased hydrocinnamic acid production by *C. sporogenes.* To confirm this, we tested a *C. sporogenes* Δ*fldH* deletion mutant^46^. The *fldH* gene encodes phenyllactate dehydrogenase, an upstream enzyme in the Phe metabolism pathway, and its deletion abrogates hydrocinnamic acid production from Phe present in the culture medium^45^. The baseline hydrocinnamic acid production of the *C. sporogenes* Δ*fldH* knockout was significantly lower compared to the wildtype control (Fig. 8C). Upon addition of Eld extract, *C. sporogenes* Δ*fldH* hydrocinnamic acid production reached levels comparable to the wildtype control after 24 hours. Taken together, these findings demonstrate that Eld extract can be used by gut microbes to yield hydrocinnamic acid, and this could contribute to the *in vivo* mechanism by which dietary Eld supplementation resulted in increased hydrocinnamic acid levels in portal plasma.

## Discussion

A healthy lifestyle can be a more accessible alternative approach to prevent and treat metabolic diseases, compared to pharmaceutical therapies. An aspect of this is maintaining a healthy diet consisting of fruits, vegetables, grains, and green tea; components which are rich in phenolic compounds such as flavonoids and other polyphenols^24,47^. While several of these compounds have been implicated in the prevention of metabolic diseases^24,28,48–50^, the exact mechanism remains unclear. It is well-established that certain members of the gut microbiome metabolize polyphenols into monophenolic acids^23^, and our group demonstrated that the gut microbial catabolite 4-HPAA is key in reversing obesity-associated hepatic steatosis in mice^28^. In the current study, we further investigate the contribution of the gut microbiome and dietary Eld supplementation on physiological parameters associated with obesity in male and female mice. We used an ABX cocktail to ablate the gut microbiota of animals on a 12-week dietary prevention study and observed microbiome-dependent improvements in insulin sensitivity and hepatic steatosis.

Our findings contribute mechanistic insights to previous observations on the insulin sensitivity improving properties of Eld^51–53^. We reveal that the dietary effect on insulin homeostasis can in part be attributed to the presence of an intact gut microbiome, thereby emphasizing the importance of bacterial metabolism for the Eld prebiotic. This Eld-microbiota effect on insulin sensitivity was comparable, if not more profound than supplementation with the well-studied MPA PCA at similar dietary dosage. This underscores the potency of hydrocinnamic acid, and implies that Eld metabolism could yield several bioactive metabolites. Notably, the microbiome-dependent effect of dietary elderberry had a stronger phenotype in female mice with respect to insulin sensitivity, while male mice exhibited a more profound improvement of their fatty liver phenotype. These findings mimic previously observed sex differences in mice and humans with metabolic diseases, as well as emphasizes the importance of including both male and female research participants^9,10,54,55^.

We developed a new LC-MS/MS method for detecting monophenolic acids, chromatographically separating structural isomers, and observed gut microbial production of hydrocinnamic acid and 3-HPAA from an Eld extract substrate, in addition to the 4-HPAA and vanillic acid catabolites we previously reported^28^. We proceeded to identify hydrocinnamic acid as a key microbial metabolite responsible for hepatic steatosis alleviation and improvements in lipid metabolism, via activation of AMPKα. It is notable that robust AMPKα activation is achieved at submicromolar, physiologically relevant concentrations of hydrocinnamic acid, similar to our previous findings with 4-HPAA^28^. In addition to its role in AMPKα activation, hydrocinnamic acid has been reported to improve glucose metabolism and regulate insulin secretion, primarily due to its activity as an agonist for pancreatic free fatty acid receptor 1, a promising anti-diabetic target^56–58^. Hydrocinnamic acid also improves mitochondria biogenesis and function in muscle tissue^59^, indicating the compound could help combat the effects of aging. Our current work uncovers a novel hydrocinnamic acid activity in reducing fatty liver in a preclinical obesity model.

Previous studies have explored flavonoid supplementation in reducing fatty liver disease^60–63^, along with bacterial metabolites derived from flavonoids, such as PCA^64–66^, 4-HPAA^28^, vanillic acid^26,27^, and gallic acid^67,68^. For our study, we investigated the role of dietary Eld in comparison to the well-studied monophenolic acid PCA as a positive control. PCA has reported activities in reducing HFD-induced obesity^25,69^, inflammation^70,71^, and fatty liver disease^48,66,72^ when it’s consumed in a typical dietary amount. Conversely, it can be toxic at high doses, especially when administered via intraperitoneal injection, as it can cause significant hepatotoxicity and nephrotoxicity associated with glutathione depletion^73^. Elderberry has been traditionally used as a supplement because of its attributed abilities to reduce oxidative stress, reduce glycemia, and stimulate the immune system^74^. Eld extract is rich in cinnamic acid^29^, as well as the flavonoids quercetin and rutin, which can be metabolized by the gut microbiota to monophenolic acids such as 3-HPAA^75,76^.

Focusing on gut microbial metabolism of the Eld extract, we observed that *C*. *sporogenes* produces hydrocinnamic acid not only from Phe metabolism via a well-characterized reductive enzymatic pathway, but also from cinnamic acid which is present in a significant amount in the Eld extract. We validated this by blocking the upstream part of the Phe to hydrocinnamic acid metabolism pathway using a *C. sporogenes* Δ*fldH* knockout strain, which lacks phenyllactate dehydrogenase (Fig. 8A). Upon incubation with Eld extract, *C. sporogenes* Δ*fldH* sustained production of hydrocinnamic acid, thereby confirming our hypothesis that hydrocinnamic acid can be produced in substantial quantities from Eld substrates like cinnamic acid. *C. sporogenes* converts cinnamic acid to hydrocinnamic acid using the enzyme acyl-CoA dehydrogenase (AcdA, Fig. 8A)^45,77 78^ as the last step in the Phe metabolism pathway.

Not only does this bypass the most energetically demanding step in Phe metabolism pathway (dehydration of phenyllactic acid to cinnamic acid)^79,80^, it also suggests that Eld provides an additional substrate for hydrocinnamic acid production under Phe-limiting conditions. Other Clostridia harbor the genes responsible for the reduction of Phe, for example *Clostridium cadaveris*, and *Peptostreptococcus anaerobius*^81^. Further experimentation using germ-free animals monocolonized with these and other candidate hydrocinnamic acid producing gut commensals will reveal whether they contribute to host metabolic health.

Our study has several limitations, including the potential effect of ABX on mouse weight gain and food intake as well as impairment of accurate gene expression in liver lysates^82^. Although these ABX-specific effects could be circumvented by using a gnotobiotic model, germ-free mice are mostly resistant to HFD-induced obesity^83^, and large multi-arm feeding studies are not feasible for practical reasons. Additionally, collecting blood plasma from fasted animals interfered with the detection of most hormones other than leptin. This collection from fasted animals might likewise prevent the LC-MS/MS detection of potentially interesting diet- and microbiota-derived metabolites with a short half-life in circulation.

In conclusion, our study underscores the importance of gut microbial metabolism in dietary prevention with a polyphenol-rich elderberry extract in the context of obesity-associated metabolic disease parameters. We identified hydrocinnamic acid as a potent microbe-derived metabolite capable of activating host AMPKα, and which is likely a key mechanistic contributor to improved insulin sensitivity and abrogation of obesity-associated hepatic steatosis. Our comparison of male and female animals revealed common effects on hepatic gene expression, AMPKα activation, gut microbial metabolism and insulin tolerance, as well as sex-specific differences in accumulation of fat in the liver. Overall, our in-depth analysis adds hydrocinnamic acid to a growing list of potent bioactive diet-microbe cometabolites which will help inform future dietary and probiotic interventions.

## Methods

### Animal studies

Male and female C57BL/6 mice were bred and maintained under specific pathogen-free (SPF) conditions within the Biological Resources Unit (BRU) of Cleveland Clinic Lerner Research Institute. At 4 weeks old, animals were given ad libitum access to a standard HFD (D12492, Research Diets, Inc.), or the same HFD supplemented with 1% w/w Elderberry extract powder (70120034, Artemis International), or 1% w/w Protocatechuic acid (PCA) (BT211395, Avantor) for 12 weeks. Mice were also allowed free access to either regular water or antibiotic cocktail (ABX) water that consists of 1 g/L neomycin (21810031, Life Technologies), 1 g/L ampicillin (BP176025, Fisher Healthcare), and 0.5 g/L vancomycin (V06500, Research Products International). All of our experiments and procedures were approved by the Institutional Animal Care and Use Committee (IACUC).

Animal food consumption and body weights were measured weekly for the entire duration of the experiment. At endpoint, lean and fat mass were measured using EchoMRI body composition analysis (EchoMRI, LLC). Endpoint plasma levels of leptin, active glucagon-like peptide 1 (GLP-1), glucagon, insulin, and peptide YY (PYY), were measured by U-PLEX Gut Hormone Combo 1 assay according to manufacturer’s instructions (Meso Scale Diagnostics, Rockville, MD, USA). Right lobes of the murine livers were fixed by submersion in formalin for 16 hours, then dehydrated in histological grade ethanol, embedded in paraffin, sectioned, and stained with hematoxylin and eosin (H&E).

### Glucose and insulin tolerance test

On week 10, mice were fasted for 4 hours and then injected intraperitoneally with 1 g/kg glucose for ITT. We measured blood glucose by using Accucheck glucometers before the glucose injection (baseline) and 15, 30, 60, and 120 min after the glucose injection. For ITT on week 11, mice were fasted for 4 hours and then injected intraperitoneally with 0.5U/kg insulin. We measured blood glucose by using Accucheck glucometers before the glucose injection (baseline) and 15, 30, 60, and 120 min after the glucose injection. During the ITT test, animals showing signs of severe hypoglycemia, observed by testing blood glucose levels, were immediately injected with glucose 1 g/kg I.P. to rescue those animals from the shock, according to the guidelines outlined in our IACUC protocol.

### Cecal microbiome sequencing and analysis

DNA from mouse cecal contents was harvested using the QIAGENPowerSoil Pro kit according to the manufacturer’s protocol. 16S rRNA gene amplicon sequencing and bioinformatics analysis were performed using methods explained earlier^84–86^. Briefly, raw 16S amplicon sequence and metadata, were demultiplexed using the split_libraries_fastq.py script implemented in QIIME2^87^. The demultiplexed fastq file was split into sample-specific fastq files using split_sequence_file_on_sample_ids.py script from QIIME2. Individual fastq files without non-biological nucleotides were processed using Divisive Amplicon Denoising Algorithm (DADA) pipeline^88^. The output of the dada2 pipeline (feature table of amplicon sequence variants (an ASV table)) was processed for alpha and beta diversity analysis using phyloseq^89^, and microbiomeSeq (http://www.github.com/umerijaz/microbiomeSeq) packages in R. We analyzed variance (ANOVA) among sample categories while measuring the of α-diversity measures using plot_anova_diversity function in microbiomeSeq package. Permutational multivariate analysis of variance (PERMANOVA) with 999 permutations was performed on all principal coordinates obtained during CCA with the *ordination* function of the microbiomeSeq package. Pairwise correlation was performed between the microbiome (genera) and metabolomics (metabolites) data was performed using the microbiomeSeq package.

### Statistical analysis

Differential abundance analysis was performed using the random-forest algorithm, implemented in the DAtest package (https://github.com/Russel88/DAtest/wiki/usage#typical-workflow). Briefly, differentially abundant methods were compared with False Discovery Rate (FDR), Area Under the (Receiver Operator) Curve (AUC), Empirical power (Power), and False Positive Rate (FPR). Based on the DAtest’s benchmarking, we selected LEfSe and ANOVA as the methods of choice to perform differential abundance analysis. We assessed the statistical significance (*P* < 0.05) throughout, and whenever necessary, we adjusted *P*-values for multiple comparisons according to the Benjamini and Hochberg method to control False Discovery Rate^90^. Linear regression (parametric test), and Wilcoxon (Non-parametric) test were performed on genera and ASVs abundances against metadata variables using their base functions in R (version 4.1.2; R Core Team^91^).

### Quantitative real-time polymerase chain reaction

10-20 mg of snap-frozen liver tissue was lysed using the tissue homogenizer in 1 mL TRIzol reagent (15596018, Thermo Fisher Scientific). Next, 200 μL chloroform was added, the sample was vortexed vigorously and centrifuged at 12,000 x *g* for 15 minutes. The upper clear layer was transferred to another tube with 500 μL isopropanol, centrifuged at 12,000 x *g* for 17 minutes until the RNA pellet precipitated. Finally, the RNA pellet was washed with 1 mL 75% ethanol and resuspended in molecular grade water. DNase treatment was performed using the DNA-*free*™ DNA Removal Kit (AM1906, Thermo Fisher Scientific) according to manufacturer instruction. cDNA was generated using qScript cDNA SuperMix (101414-106, Quanta-Bio) using ∼1 µg of RNA as template. Real-time PCR was performed using an Applied Biosystems Step One Plus system. mRNA expression levels were calculated based on the ΔΔ-CT method and genes of interest were normalized to the house keeping gene *CycloA*. A list of primer sequences can be found in Table 1.

**Table 1:**
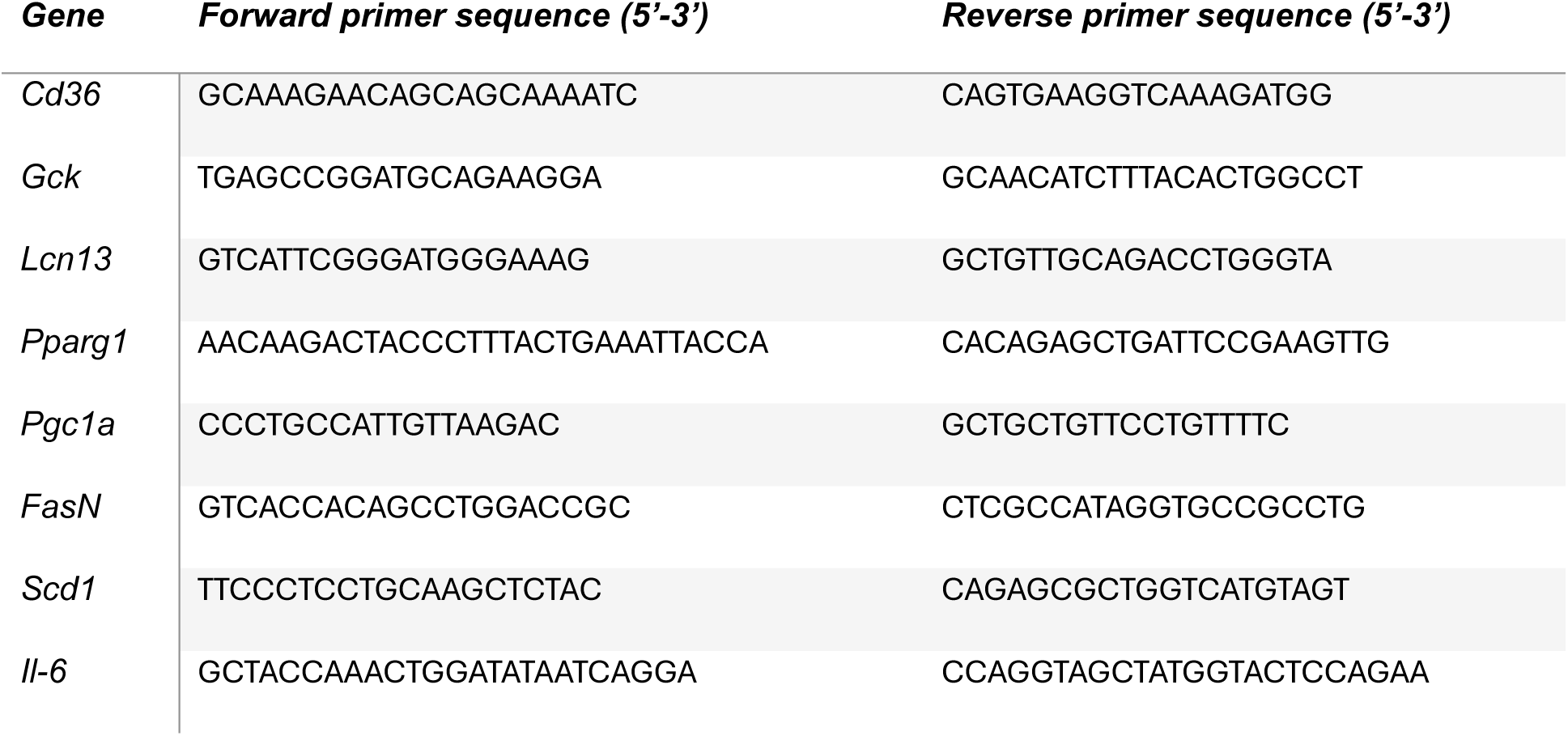
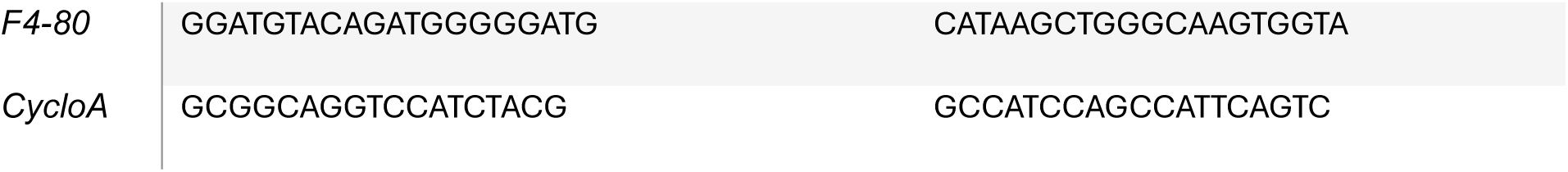
Primer sequences for quantitative real-time polymerase chain reaction.

### LC-MS/MS analysis of monophenolic acids

Portal plasma or bacterial supernatant were prepared for LC-MS/MS analysis and quantified as previously mentioned^92^. Stable-isotope-dilution high performance liquid chromatography tandem mass spectrometry (LC–MS/MS) was used for quantification of levels of different phenolic acids using a LCMS-8050 Triple Quadrupole (Shimadzu Scientific Instruments, Inc., Columbia, MD, USA), equipped with an electrospray ionization source operating in negative ion mode. XSelect HSS T3 Column (4.6 mm X 250 mm; 3.5 µm) (Cat # 186004788, Waters) was used for chromatographic separation. The D_6_ 4-HPAA was used as an internal standard (D-7842, CDN Isotopes) at 10 μM. Monophenolic acids were monitored using multiple reaction monitoring (MRM) of precursor and characteristic product ions in negative mode as follows: *m/z* 157.1 → 112.9 for D6-4-HPAA; *m/z* 149 → 105.05 for hydrocinnamic acid (3-phenylpropionic acid); *m/z* 151.0 → 106.9 for 3-hydroxyphenyl acetic acid; *m/z* 193.0 → 178.15 ferulic acid; *m/z* 137.0 → 92.9 for 4-hydroxybenzoic acid (4-HBA); *m/z* 153.2 → 108.9 for protocatechuic acid (PCA); *m/z* 167 → 152.1 for vanillic acid (VA); *m/z* 165.1 → 121.0 for phloretic acid (PA); *m/z* 150.9 → 106.9 for 4-hydroxyphenyl acetic acid; *m/z* 151.2 → 107 for 2-hydroxyphenyl acetic acid.

### Cell culture and immunoblotting

Human hepatoma cells (HuH-7) were seeded at a density of 1X10^6^/well in 6-well tissue culture plates (TP92406, MIDSCI) in Dulbecco’s Modified Eagle Medium (DMEM) containing 10% fetal bovine serum. Media was changed two hours prior to treatment with a DMSO vehicle control, hydrocinnamic acid (AAA1490822, Thermo Scientific Chemicals), or 3-HPAA (AAA1490822, Thermo Scientific Chemicals) for 60 minutes. Cell pellets or murine liver homogenates were suspended in radioimmunoprecipitation assay (RIPA) buffer with protease and phosphatase inhibitors (Thermo Scientific).

Protein concentration was quantified using a Bradford Assay kit according to the manufacturer’s instructions (Thermo Scientific). Gel electrophoresis was carried out with samples diluted in Laemmli buffer with 2-mercaptoethanol. A 12% Mini-PROTEAN® TGX™ precast gel was loaded with 10 µg of sample per well and run at 100V for 2hrs (Biorad). Proteins were transferred to a 0.2 µm PVDF membrane (Biorad) using wet electroblotting at 4°C for 16hours. Images were taken using a GE Amersham Imager 6000 and ImageJ^93^ was used for densitometric protein expression analysis. Primary antibodies (pT172-AMPKα, 2535, Cell Signaling), (AMPKα, 5831, Cell Signaling), (β-actin, C825B47, Proteintech) were prepared 1:1000 in TBST buffer with 5% BSA (w/v) (pS79-ACC, 11818, Cell Signaling), (ACC, 3676, Cell Signaling), were prepared 1:10000 in TBST buffer with 5% (w/v) nonfat dried milk. Rabbit secondary (7074, Cell Signaling) and mouse secondary (405306, BioLegend) were prepared 1:5000 in TBST buffer with 5% BSA.

### Microbial culturing

Microbial culture experiments were performed in an anaerobic chamber (Coy Laboratory Products, Inc.) under the following conditions: 90% N_2_, 5% H_2_, 5% CO_2_, and < 25 ppm O_2_. To examine the conversion of Eld into hydrocinnamic acid, *C. sporogenes* ATCC 15579, *C. sporogenes* Δ*fldH*Δ*cutC*, and *B. fragilis* 3_1_12 HM-20 were plated on a 50:50 (v/v) mixture of Wilkins-Chalgren and Gifu Anaerobic or Tryptic Soy agar. Single colonies were cultured in liquid medium and incubated at 37° C for 24 hours. Next, these cultures were diluted 1:10 in fresh media with vehicle or 1% Eld extract. After 4 hours and 24 hours, bacteria were pelleted by centrifugation at 8000 x *g* for 5 minutes, and supernatant was filtered using Sterile Syringe Filters 0.22μm pore sizes (SLGVM33RS, MilliporeSigma). Next, the filtered supernatant was prepped and analyzed by LC-MS/MS as described above.

## Data availability statement

All data are available in the main text or the supplementary materials. Raw sequence files from the 16S rRNA gene sequencing were deposited in Zenodo’s Sequence Read Archive (accession DOI: 10.5281/zenodo.15208086).

## Supporting information

Supplemental Images

## Acknowledgements

The authors would like to thank Artemis International for providing the Eld extract. We thank Stanley Hazen for providing us the *C. sporogenes* Δ*fldH*Δ*cutC* strain. This work was supported by seed funding from the Cleveland Clinic Foundation (J.C.), and in part by a Research Grant from the Prevent Cancer Foundation (PCF2019-JC), and a National Institutes of Health (NIH) grant R01 AI153173 (J.C.). I.N. was in part supported by the NIH R01HL16074 grant.

## Author contributions

S.A. and J.C. conceived and designed the experiments. S.A., B.D., J.C., L.J.O., R.L.M., V.B., T.T., and L.H.N performed the mouse experiments. S.A. performed the biochemical workup of mouse experiments. S.A., and S.P. performed and analyzed the *in vitro* microbial Eld catabolism experiments. N.S. conducted the microbial sequencing analysis. S.A., and I.N. developed the LC-MS/MS method and performed subsequent analysis of *in vivo* and *in vitro* samples. C.G.F. performed the histological analysis of murine liver sections. S.A. and J.C. designed experiments and or analyzed the subsequent data. S.A. and J.C. wrote the manuscript and handled visualization. All authors discussed the results and commented on the manuscript.

## Competing interest declaration

J.C. was a Scientific Advisor for Seed Health, Inc. The other authors declare they have no competing interests.

