## Supplemental Images for "Gut microbial conversion of dietary elderberry extract to hydrocinnamic acid improves obesity-associated metabolic disorders"

**This PDF file includes:**
Figs. S1 to S7

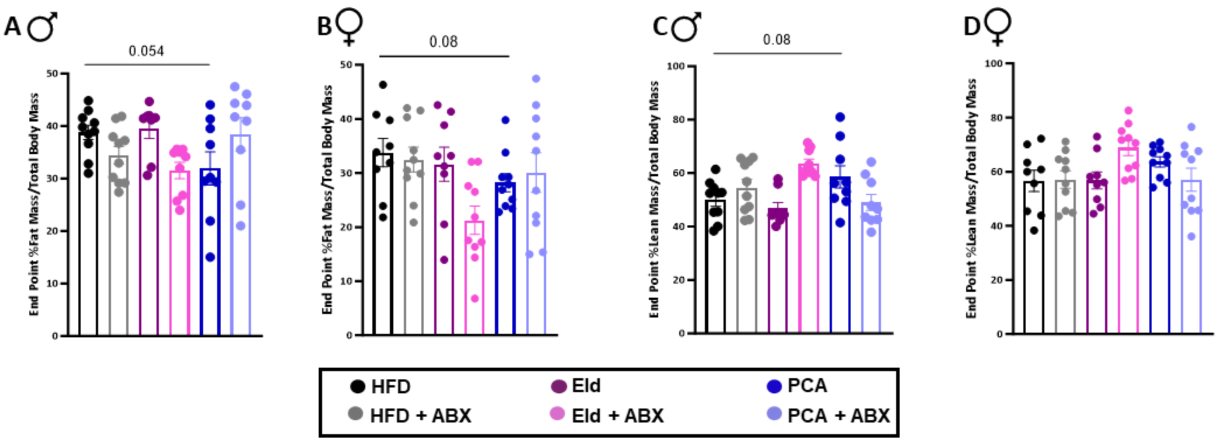


**SI Fig. S1. Body composition (%lean mass and %fat mass) between the different groups in both males and females. (A-B)** Fat mass, and **(C-D)** Lean mass for male and female mice after 12 wks on diet as measured by EchoMRI and normalized to total body mass, n = 9–10 per group. Error bars represent SEM. Statistical analysis was performed using unpaired two-tailed Student's t test.

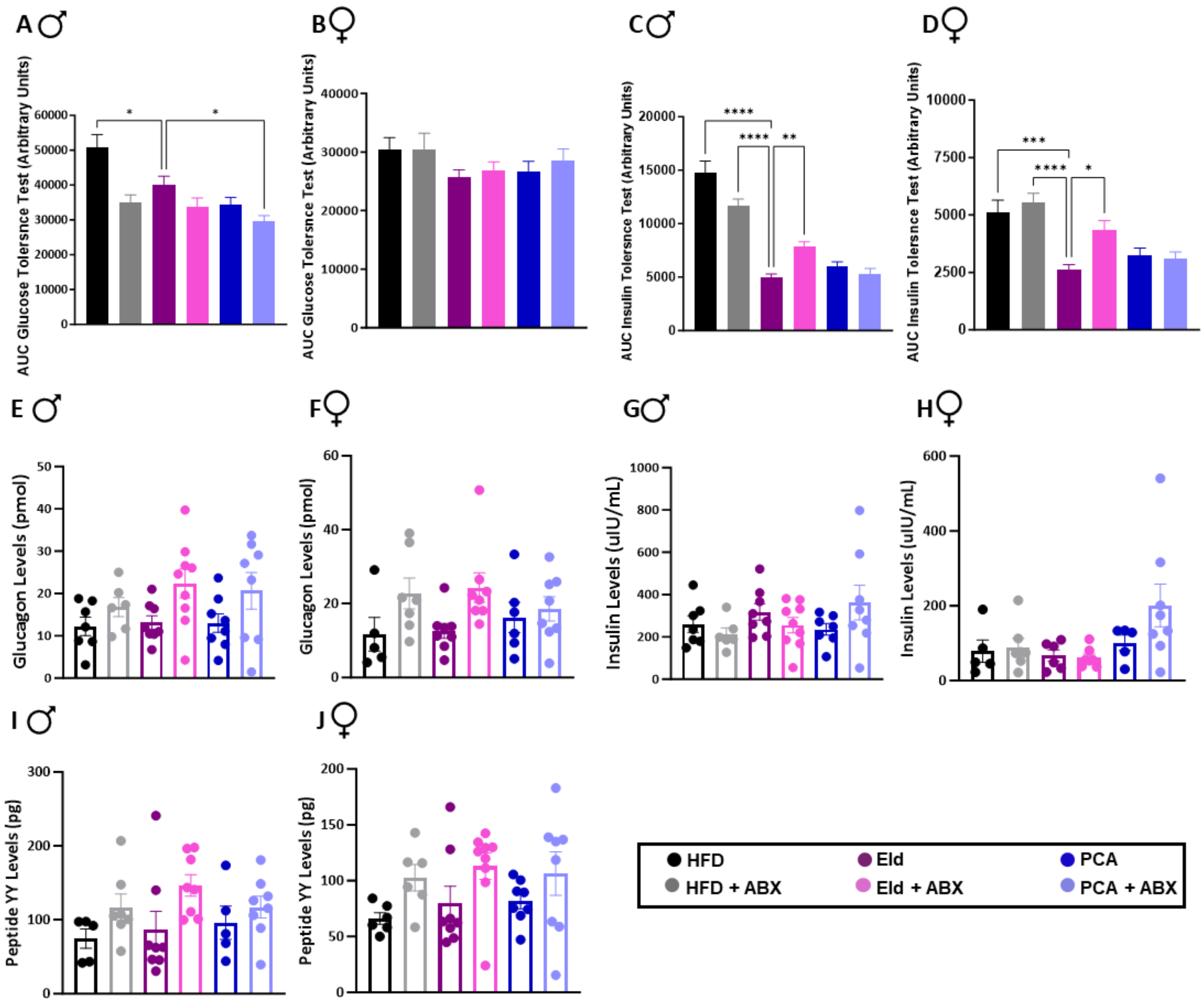


**SI Fig. S2. Area under the curve for glucose tolerance test and insulin tolerance test, glucagon, insulin, and peptide YY levels in the end point plasma of both males and females. (A-B)** Mean cumulative area under the curve (AUC) for male and female mice glucose tolerance test on week 10. **(C-D)** Mean cumulative area under the curve (AUC) for male and female mice insulin tolerance test on week 11, error bars represent SEM. Statistical analysis was performed via ANOVA. End point levels of male and female plasma **(E-F)** glucagon, **(G-H)** insulin, and **(I-J)** peptide YY. Error bars represent SEM. Statistical analysis was performed via ANOVA.

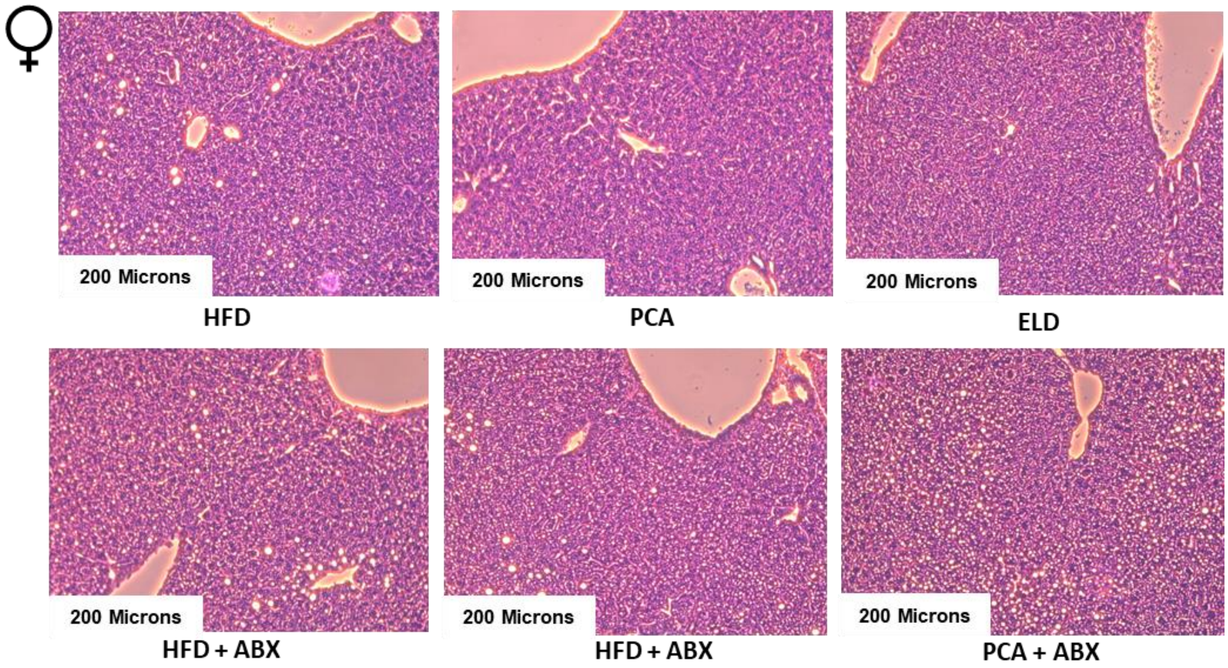


**SI Fig. S3. H&E-stained livers of female mice on the different diets.** After 12 wks of the different diets, H&E-stained sections of the female mouse livers. Representative images shown from n = 6 per group.

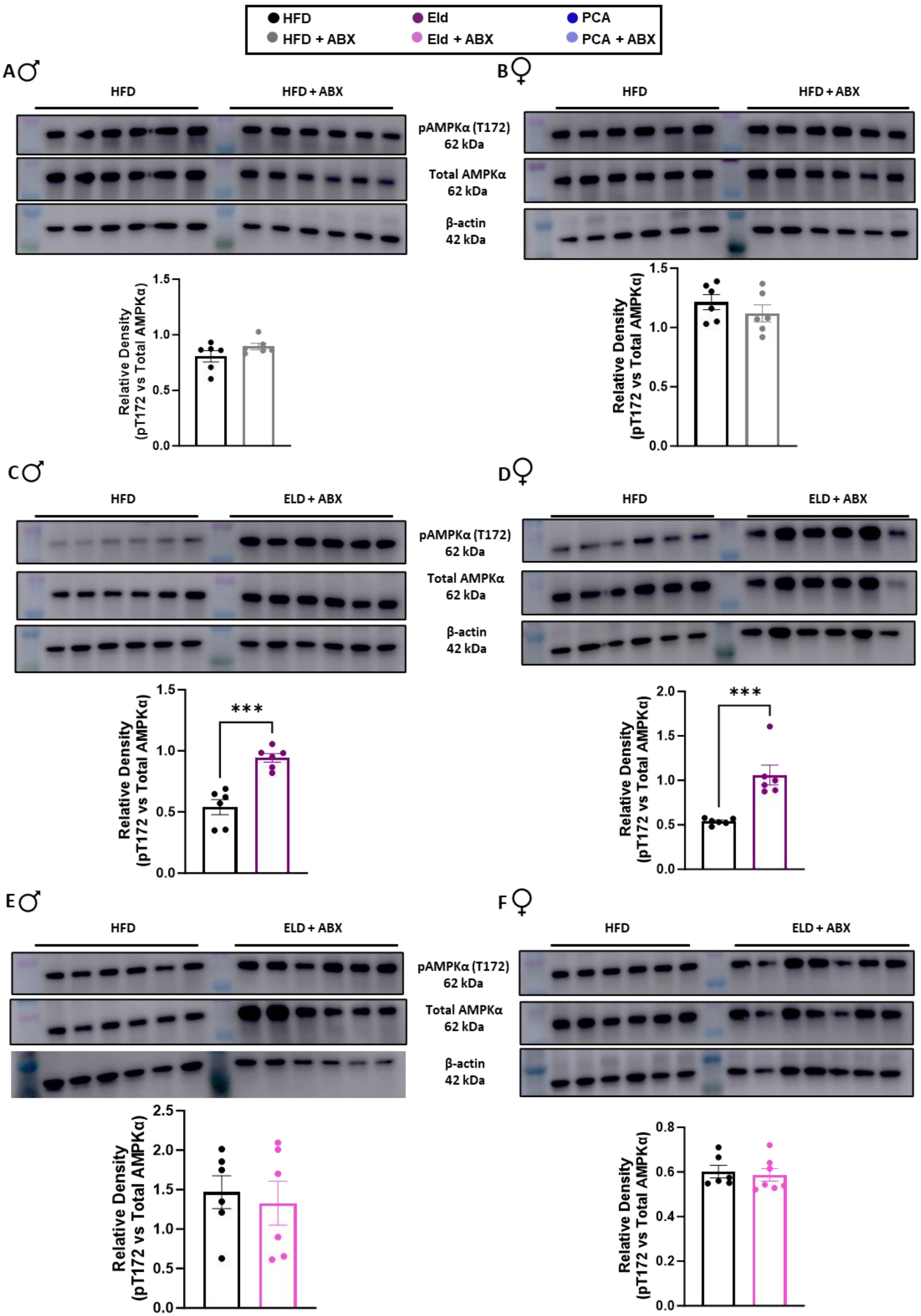

HFD
Eld
PCA
HFD + ABX
Eld + ABX
PCA + ABX
A♂
B♀
C♂
D♀
E♂
F♀
HFD
HFD + ABX
ELD + ABX
pAMPKα (T172)
62 kDa
Total AMPKα
62 kDa
β-actin
42 kDa
Relative Density
(pT172 vs Total AMPKα)
***

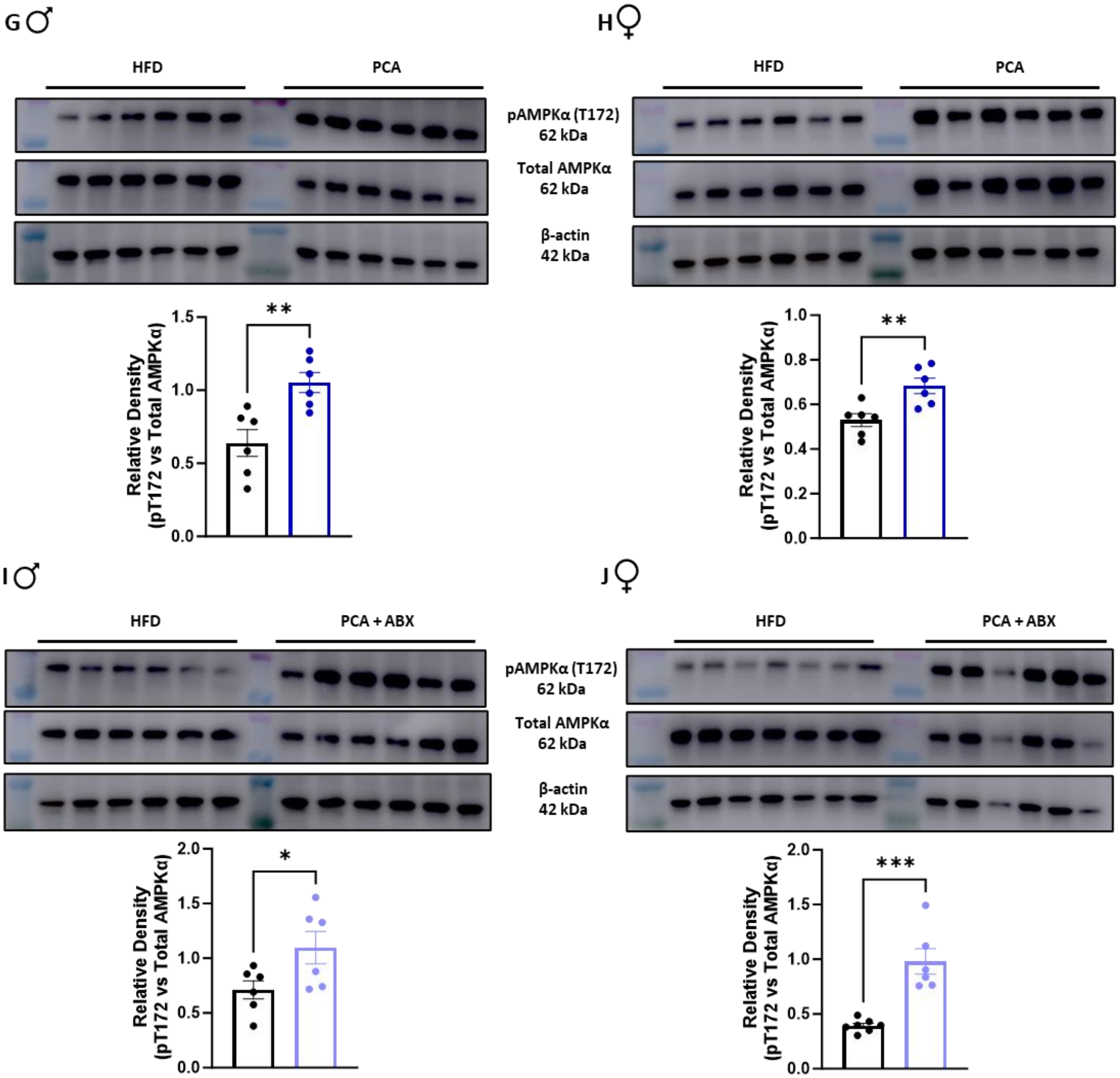


**SI Fig. S4. Activation of AMPKα in male and female mice in each group compared to the high fat diet control group. (A)** Top: Western blot analysis of pAMPKα (T172), total AMPKα, and β-actin in male mice on high fat diet and antibiotic water compared to mice high fat diet and regular water after 12 wks. Bottom: densitometric quantification of the membrane, n = 6 per group **(B)** Female mice. **(C)** Top: Western blot analysis of pAMPKα (T172), total AMPKα, and β-actin in male mice on high fat diet with 1% elderberry extract and regular water compared to mice high fat diet and regular water after 12 wks. Bottom: densitometric quantification of the membrane, n = 6 per group **(D)** Female mice. **(E)** Top: Western blot analysis of pAMPKα (T172), total AMPKα, and β-actin in male mice on high fat diet with 1% elderberry extract and antibiotic water compared to mice high fat diet and regular water after 12 wks. Bottom: densitometric quantification of the membrane, n = 6-7 per group **(F)** Female mice. **(G)** Top: Western blot analysis of pAMPKα (T172), total AMPKα, and β-actin in male mice on high fat diet with 1% PCA and regular water compared to mice high fat diet and regular water after 12 wks. Bottom: densitometric quantification of the membrane, n = 6 per group **(H)** Female mice. **(I)** Top: Western blot analysis of pAMPKα (T172), total AMPKα, and β-actin in male mice on high fat diet with 1% PCA and antibiotic water compared to mice high fat diet and regular water after 12 wks. Bottom: densitometric quantification of the membrane, n = 6-7 per group **(J)** Female mice. Error bars represent SEM. Statistical analysis was performed using unpaired two-tailed Student's t test.

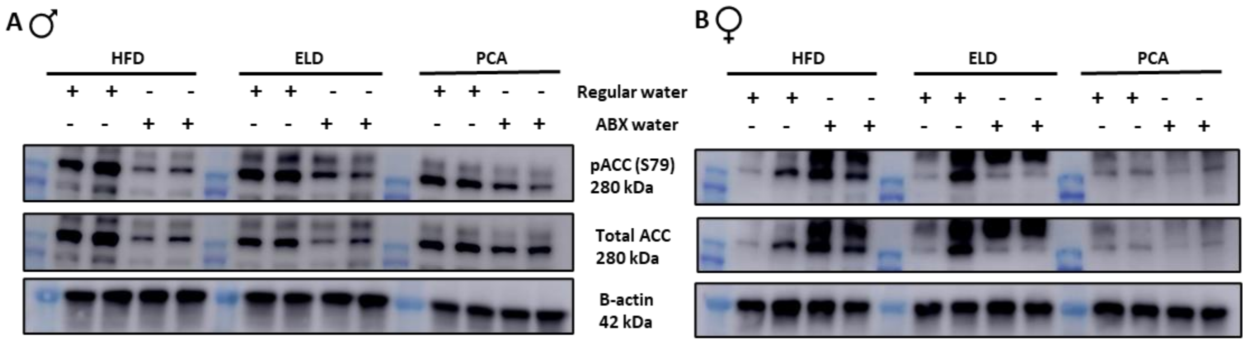


**SI Fig. S5. Acetyl-CoA Carboxylase activation in the end livers of both males and females. (A-B)** Western blot analysis of pACC (S79), total ACC, and β-actin in male and female mice after 12 wks of the different diets and water.

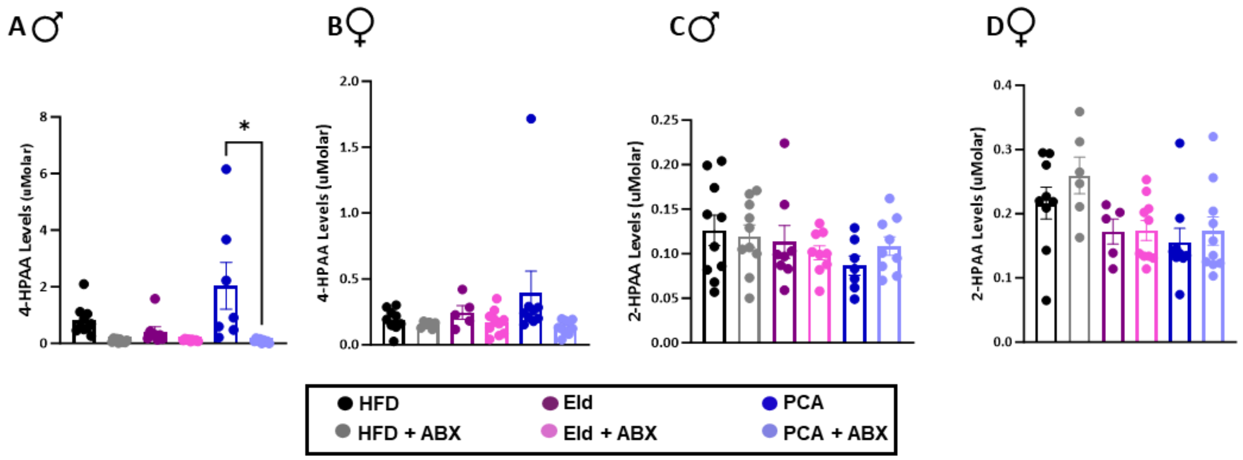


**SI Fig. S6. No difference in 4-HPAA and 2HPAA levels in the end point plasma of both males and females. (A-B)** 4-Hydroxyphenyl acetic acid (4-HPAA), **(C-D)** 2-Hydroxyphenyl acetic acid (2-HPAA) LC-MS/MS measured Portal plasma levels from male and female mice (n = 5–9 per group). Error bars represent SEM. Statistical analysis was performed with one-way ANOVA.

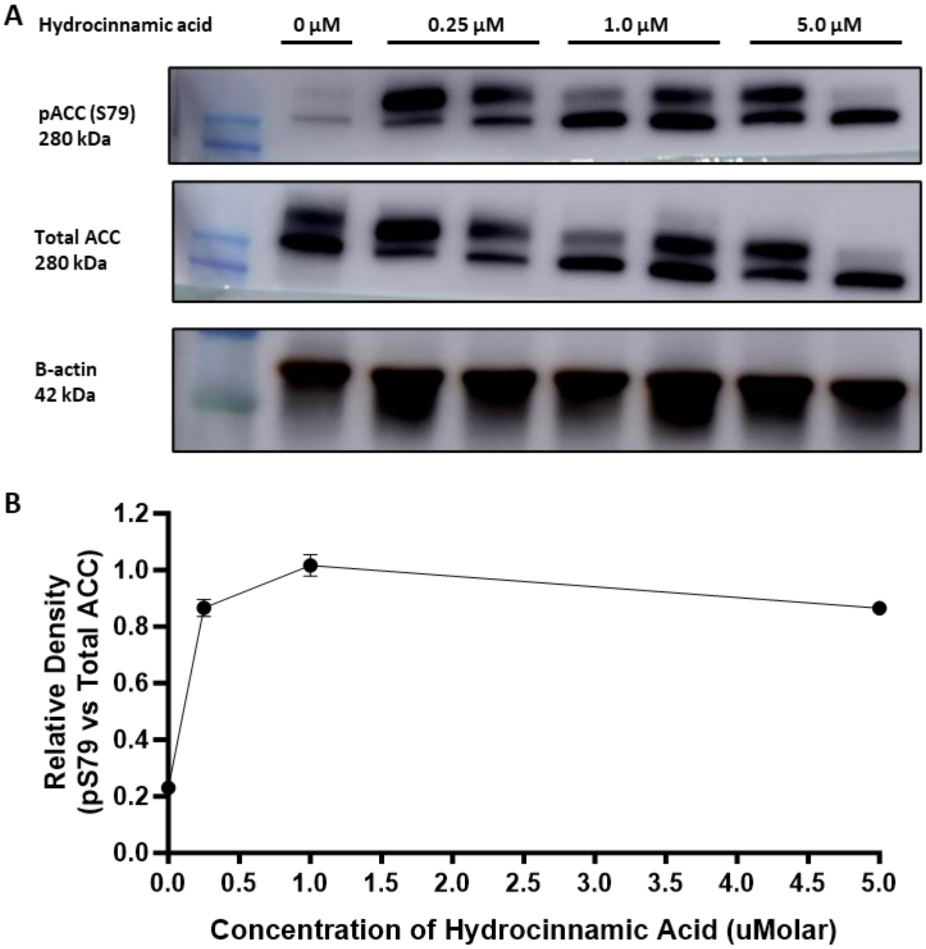


**SI Fig. S7. Hydrocinnamic acid inactivates Hepatic in acetyl-CoA carboxylase (ACC) in HuH-7 cells in a physiological level. (A)** Western blot analysis of pACC (S79), total ACC, and β-actin for HuH-7 cells after treatment with either DMSO vehicle, 50 µM 3-PPA, or 50 µM hydrocinnamic acid. **(B)** Relative expression of the ratio of ACC activation after treatment with different physiological concentrations of hydrocinnamic acid. Error bars represent SEM.
